# A data-driven approach to establishing cell motility patterns as predictors of macrophage subtypes and their relation to cell morphology

**DOI:** 10.1101/2022.11.29.518400

**Authors:** Manasa Kesapragada, Yao-Hui Sun, Kan Zhu, Cynthia Recendez, Daniel Fregoso, Hsin-Ya Yang, Marco Rolandi, Rivkah Isseroff, Min Zhao, Marcella Gomez

## Abstract

The motility of macrophages in response to microenvironment stimuli is a hallmark of innate immunity, where macrophages play pro-inflammatory or pro-reparatory roles depending on their activation status during wound healing. Cell size and shape have been informative in defining macrophage subtypes. Studies show pro and anti-inflammatory macrophages exhibit distinct migratory behaviors, in vitro, in 3D and in vivo but this link has not been rigorously studied. We apply both morphology and motility-based image processing approaches to analyze live cell images consisting of macrophage phenotypes. Macrophage subtypes are differentiated from primary murine bone marrow derived macrophages using a potent lipopolysaccharide (LPS) or cytokine interleukin-4 (IL-4). We show that morphology is tightly linked to motility, which leads to our hypothesis that motility analysis could be used alone or in conjunction with morphological features for improved prediction of macrophage subtypes. We train a support vector machine (SVM) classifier to predict macrophage subtypes based on morphology alone, motility alone, and both morphology and motility combined. We show that motility has comparable predictive capabilities as morphology. However, using both measures can enhance predictive capabilities. While motility and morphological features can be individually ambiguous identifiers, together they provide significantly improved prediction accuracies (75%) from a training dataset of 1000 cells tracked over time using only phase contrast time-lapse microscopy. Thus, the approach combining cell motility and cell morphology information can lead to methods that accurately assess functionally diverse macrophage phenotypes quickly and efficiently. This can support the development of cost efficient and high through-put methods for screening biochemicals targeting macrophage polarization.

**Author summary:** Previous work has shown that macrophage phenotypes can be distinguished by their morphological characteristics. We extend this work to show that distinct motility patterns are linked to macrophage morphology. Thus, motility patterns can be used to differentiate phenotypes. This can enable high-throughput classification of cell phenotypes without regard for the high-resolution images needed to quantify morphological characteristics. Furthermore, combining motility-based features with morphological information improves prediction of macrophage subtypes by a machine learning based classification model.

## Introduction

Macrophages reside in or are recruited to all tissues in the body. They play an important role in tissue hemostasis and are an essential part of the human defense system. In wound healing, they are among the first cell types to traffic to the wound. They form an immune defense line, promote, and resolve inflammation, remove dead cells, and cell debris, and support cell proliferation and tissue restructure [1, 2]. There exists a continuum of macrophage functions [3, 4], which have been historically binned into two categories with primarily pro-inflammatory and anti-inflammatory functions, respectively. The two major groups of macrophages that display different functional phenotypes present in the wound bed are referred to as classically activated (pro-inflammatory) and alternatively activated (anti-inflammatory) macrophages. During the inflammation phase, several mediators such as bacterial products like lipopolysaccharide (LPS) and inflammatory cytokines like interferon (IFN)-*γ* can stimulate macrophages to differentiate into pro-inflammatory status [5, 6]. Pro-inflammatory macrophages exhibit antimicrobial properties through the release of inflammatory mediators inducing tumor necrosis factor (TNF)-*α*, IL-1*β*, nitric oxide (NO) and interleukin (IL)-6 [7]. Prolonged persistence of pro-inflammatory macrophages without transitioning to anti-inflammatory phenotypes, however, is rather detrimental to wound tissues, which will likely stall, or delay wound healing [8, 9]. On the other hand, macrophages activated by IL-4 and IL-13 develop into so-called alternatively activated macrophages (anti-inflammatory) that suppress inflammatory reactions and adaptive immune responses and play a reparative role during the proliferation and maturation phases [10]. However, like pro-inflammatory macrophages, if anti-inflammatory macrophages persist for too long hypertrophic scars and keloids develop due to excessive collagen production [11, 12].

The involvement of macrophage phenotypes in the wound healing process indicates that detecting macrophage status could be diagnostically useful. However, advances in the field have been hampered, largely due to lack of definitive and optimal biomarkers for macrophage phenotypes. Traditionally, characterization of pro and anti-inflammatory subtypes is carried out by quantification of multiple cell surface markers, transcription factors and cytokine profiles. These approaches are time-consuming, require large numbers of cells/tissues and are resource intensive. Difficulty detecting pro and anti-inflammatory macrophage phenotypes in vivo is compounded by similar problems in in vitro-derived macrophages. For example, Arginase-1, which is considered a classic marker for anti-inflammatory macrophages, is also upregulated in pro-inflammatory macrophages [13, 14] and protein expression of Arginase-1 or CD206 is too low for reliable flow cytometry detection [4, 5, 11, 15]. In conclusion, a more robust discriminating system is critically needed to improve the detection and understanding of macrophage phenotypes and their relation to their function.

Because of these distinct roles, macrophages undergo a precisely regulated dynamic transition in their functionalities in time and space. The ability to precisely regulate macrophage distribution, migration, and function polarization (i.e., pro vs. anti-inflammatory macrophages) that spatial-temporally corresponds to wound status offers a powerful approach to regulate wound healing. The immune response involves temporal tracking of the migration of different immune cell types in the wound. When tracking wound healing, we can use morphology and motility to track different cell types and train an algorithm to look for patterns in cellular responses characterized by identifiers, such as location (e.g., near blood vessels) or nearest cell type neighbors.

Macrophages undergo morphological changes when undergoing a change in phenotype [2]. Studies have shown that cell shape can help control macrophages — for instance, elongating them promotes behavior that enhances healing [1]. While cell size and shape have been used to characterize macrophage subtypes, how motility properties are contributing to such quantifications are not well studied, despite pro and anti-inflammatory macrophages exhibiting distinct migratory behaviors.

Machine learning methods have been successfully applied to automate the classification of different cell types from microscopy images [16, 17] and are viewed as a promising approach to high-through put screening. However, these methods are sensitive to the size of the data set, quality of the images, and sometimes facilitated by fluorescent dyes or reporters [18]. Previous work towards classifying macrophage subtype using machine learning leverages fluorescent dyes [19, 20]. Being able to identify cell types from phase contrast microscopy would simplify and accelerate macrophage classification. Furthermore, being able to identify cells based on their motility parameters can eliminate the need for high-quality imaging and result in methods robust to blurred images. We suspect that cell motility like cell morphology is linked to function. Cell tracking can be achieved with image processing techniques in real-time [21]. Motility patterns can be extracted from cell trajectories and machine learning used to classify cells based on their dynamic behavior. Finally, combining cell motility and morphological features can help elucidate the link between cell migratory behavior, morphology, and cellular phenotype.

Here we apply machine learning methods to classify macrophage subtypes and show a tight relationship between cell morphology and its motile behavior. We begin by first validating that our derived macrophage subtypes do present distinct morphological characteristics as has been shown in previous work. Morphological analysis of the label-free collective dataset suggests macrophage shapes are described by three principal shape modes. Clustering analysis is applied to identify the cells belonging to each of these three morphological groupings. Comparison with the labeled data confirms a correlation between macrophage subtype and morphology. Next, we show that independent of cell labels, there is a strong relationship between morphology and motile behavior. Motility features are computed within each cluster showing that distinct morphological features map to distinct cell motility parameters. We then apply discriminant analysis to confirm that this implies a relationship between the labeled cells with respective phenotypes and motility parameters, thereby, supporting the hypothesis that we can predict macrophage subtypes from motility behavior alone. Finally, to predict macrophage subtypes with machine learning, we propose a SVM classification model. We build multiple predictive models using morphological and motility parameters alone and in combination to predict the macrophage subtypes. A high-level schematic of the analysis carried out in this work is shown in Fig 1. We demonstrate predictions using motility are comparable to those using morphology and are equally good for the same subtype. This suggests that while they are tightly linked, they may in fact provide complementary information. We show that a combined use of motility and morphology shows an increased prediction accuracy in identifying macrophage subtypes. Hence, by combining both cell motility and morphological information we show that we can reliably detect non-activated naïve macrophages, or macrophages activated by either lipopolysaccharide (LPS) or interleukin 4 (IL-4) exposure at single-cell level.

**Fig 1.**
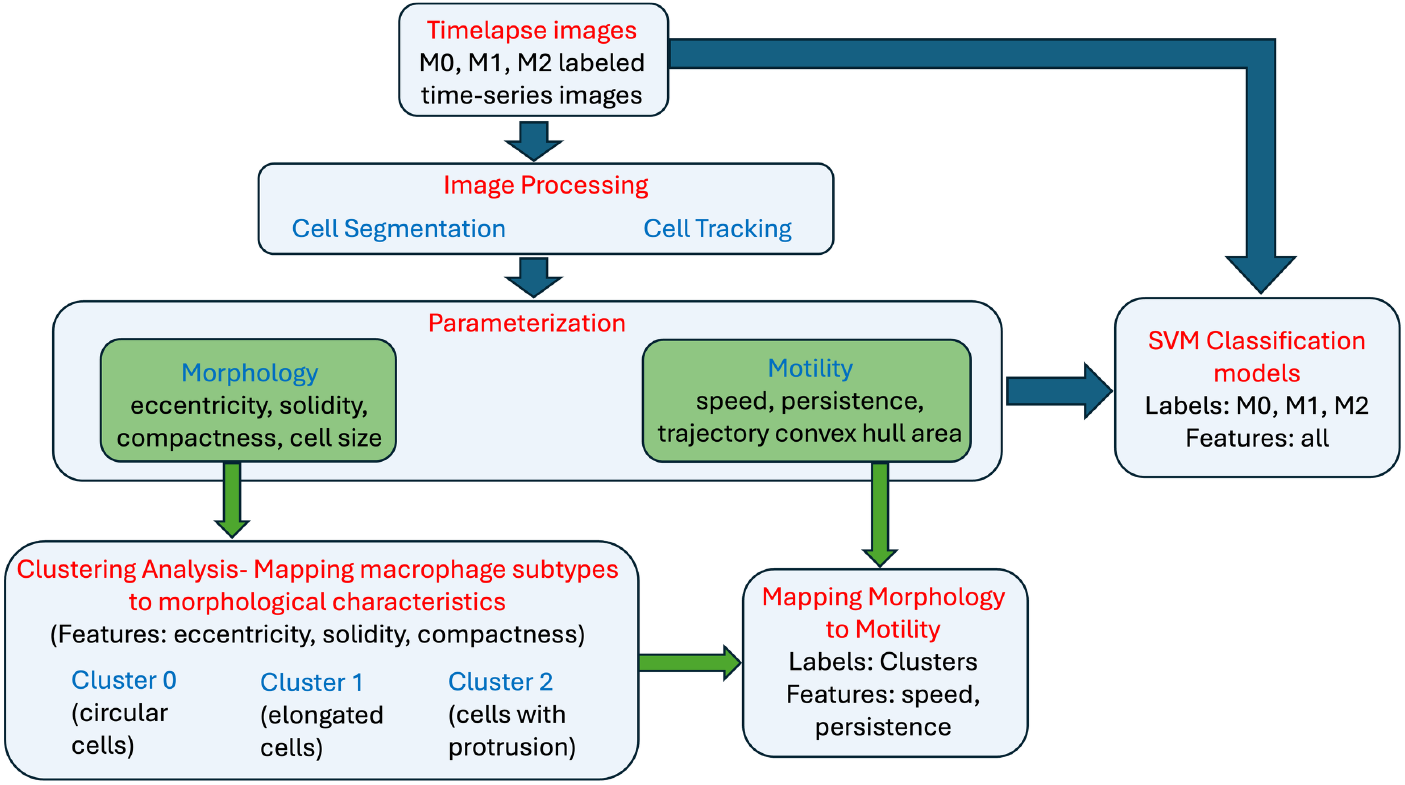
Analysis Workflow. A high-level schematic of the analysis carried out in this paper. The first step of the workflow consists of image processing methods to perform cell segmentation and cell tracking. Cell segmentation allows for parameterization of the cell morphology and cell tracking allows for parameterization of cell motility over time. Unsupervised clustering analysis on morphological features alone results in cell groupings consistent with parent image labels. Next, it is confirmed that cell motility characteristics are shared within each morphological-based cell grouping and distinct across clusters. Finally, various SVM models are presented using morphology and motility features independently and together to classify macrophage subtypes. The ground truth is provided by the parent label image.

## Materials and methods

### Isolation and culture of bone marrow-derived macrophages

The mouse strains used in this study were of the C57BL/6 background, and both male and female mice were included in the experiments. Mice were procured from Jackson Labs and housed under a strict 12-hour light cycle with a regular chow diet in a specific pathogen-free facility at the University of California (UC), Davis. All animal experiments adhered to regulatory guidelines and standards set by the Institutional Animal Care and Use Committee of UC Davis (Protocol # 21531). Bone marrow-derived macrophages (BMDMs) were generated using standard procedures as previously described [22]. Mice were euthanized using a CO2 chamber and then subjected to cervical dislocation to ensure minimal suffering. The femurs were aseptically dissected, and the bone marrow was flushed with cold Dulbecco’s phosphate-buffered saline without calcium and magnesium (DPBS). Given the endpoint nature of these experiments, no anesthesia or analgesia was applied. Bone marrow from 3-6 mice was pooled per batch to account for biological variability. The harvested bone marrow cells were cultured in DMEM (Invitrogen) supplemented with 10% Fetal Bovine Serum (Invitrogen), 1× Antibiotic-Antimycotic solution (Invitrogen), and 20% L-929 conditioned medium for six days, with an additional feeding on day 3, followed by a one-day culture without the conditioned medium. Adherent macrophages were harvested by gently scraping with a “policeman” cell scraper and used in subsequent experiments as needed. Cell viability was determined using trypan blue staining and counting. This study utilized at least four batches of BMDMs, derived from a total of 20 mice of mixed gender, generated at different times.

### Activation of bone marrow-derived macrophages

In each experiment, bone marrow-derived macrophages (BMDMs) were seeded into six tissue culture-treated well plates at varying densities and cultured in RPMI-1640 medium (Invitrogen) supplemented with 10% Fetal Bovine Serum (FBS) (Invitrogen) and 1× Antibiotic-Antimycotic solution (Invitrogen) overnight. For M1 activation, 100 ng/ml lipopolysaccharide (LPS) (Sigma, Cat number: L6143) was added to the culture medium, while for M2 activation, 20 ng/ml recombinant mouse interleukin-4 (IL-4) (R&D Systems, Cat number: 404-ML) was used [23]. Two days post-stimulation, activated M1 and M2 macrophages were employed for morphological and motility characterizations as well as functional studies. Macrophages that did not receive any stimulation served as M0 controls.

### Time-lapse recording

In each experiment, freshly differentiated macrophages were seeded in 24-well tissue culture-treated plates, cultured, and activated following the procedures described above. Forty-eight hours after stimulation, cells were monitored using a Carl Zeiss Observer Z1 inverted microscope outfitted with a motorized stage and an incubation chamber (3°C and 5% CO2). Time-lapse contrast images were captured using the MetaMorph program (Molecular Devices) with a Retiga R6 (QImaging) scientific CCD camera (6 million, 4.54 µm pixels). Typically, in each experiment, four fields of each condition were selected from different wells under a long-distance 20× objective lens. Images were taken at 5-minute intervals for up to 3 hours unless stated otherwise. We note that in some experiments, certain wells were designated for control purposes or other specific experimental requirements, and thus, they were excluded from the imaging process. In practice, to ensure the quality of our data, we exclude out-of-focus images, retaining only the most suitable ones for subsequent morphological analysis.

### Cell segmentation and tracking

The input images are segmented and tracked in a semi-autonomous fashion using the Baxter Algorithms software package [24, 25] written in Matlab. This program segments the cells and tracks them throughout each frame. It computes the 2D coordinates of a cell in each frame along with its region properties.

### Computing speed and persistence

The speed and persistence are computed at each using the following formula: The speed of a cell from trajectory points (*x*_1_, *y*_1_) to (*x*_*n*_, *y*_*n*_) where *n* is the total length of time frames, could be any point from 2 to last time frame point, *d* is the total distance travelled, *t* is the time travelled given by:

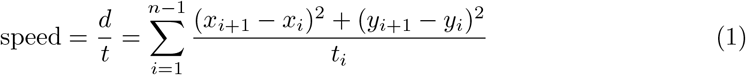

Persistence is a measure to quantify the length of a cell’s path along its entire time to know if a cell has remained closer to its initial time frame trajectory point or if it has moved farther away. It is defined as the ratio of the shortest distance traveled *p* between the initial point (*x*_1_, *y*_1_) and the *n*th time frame point (*x*_*n*_, *y*_*n*_) to the total distance traveled by the cell *d* given by:

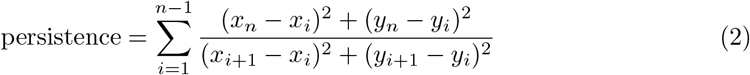

## Results

### Generation and polarization of primary murine macrophages

Bone marrow-derived macrophages (BMDMs) are primary macrophages that are differentiated from bone marrow cells [26, 27]. Murine BMDMs present an excellent model and are widely used as prototypical macrophages for investigation of mammalian macrophage functions in vitro. We isolate bone marrow from freshly cut femur bones of C57/BL6 mice. Four batches of bone marrow-derived macrophages were generated, each batch consisting of a mixture of bone marrow from 3-6 mice. In total, bone marrow from 20 mice was used for these experiments. In each experiment we grow bone marrow cells in the presence of murine macrophage colony-stimulating factor (M-CSF), as provided by supplementary L929-conditioned medium [28]. Within 6 days, the bone marrow monocyte progenitors proliferate and differentiate into a homogenous population of mature macrophages (M0). Subsequently, naïve M0 macrophages are further polarized into pro-inflammatory or anti-inflammatory macrophages by addition of bacterial lipopolysaccharide (LPS) or murine interleukin 4 (IL-4), respectively [23] (Fig. S1A). To simplify notation in the manuscript, we refer to pro-inflammatory macrophages as M1 and anti-inflammatory macrophages as M2. See the methods for details on the protocols used to differentiate macrophages into their respective phenotypes and the supplementary information for results of validation assays confirming three distinct phenotypes based on the literature. Validation assays include phagocytic capacity and quantitative PCR to determine the relative mRNA expression of a panel of selected target genes and transcription factors (see Table S2 for target genes and primers used in q-PCR).

### Generation of time-series data capturing spontaneous cell migratory patterns

Using time-lapse microscopy, we generate multiple sets of M0, M1, and M2 images, named after their most representative macrophage subtypes, taken at 5-minute intervals for up to 3 hours (Fig. S2). These are the phase contrast images without ground truths or cell labels except the image names based on culturing conditions. Although the macrophages are traditionally binned into three subtypes, they exist on a continuous spectrum, i.e., some cells may be “in-between” identified subtypes, hence we note that the parent images may contain cells of the other morphological subtypes with respect to their labels. In the current analysis, we look mainly for the dominant representations. Cell segmentation and tracking details can be found in the methods section. After cell segmentation and tracking, we select the cells that remain in the image dataset’s time-lapse for at least 1 hour. This action allows us to eliminate cells that exit the image area within less than 1 hour of their lifespan along the time series. For the characterization of cell subtypes, we compute cell morphology parameters using the first-time step and determine motility parameters based on time-lapse data. Following this, our time-lapse image datasets resulted in the identification of a total of 1018 distinct cells. These cells comprised 251 cells from M0 images, 369 cells from M1 images, and 398 cells from M2 images.

### Three principal shape modes characterize observed macrophage morphologies

We extracted and aligned [29] label-free phase contrast images of a mixed population of live macrophages. To identify the existing principal shape modes, we consider each cell in every frame of the time-lapse datasets as individual data points (n = 2329 cells). We obtain the shape information for M0 (n = 965 cells), M1 (n = 698 cells) and M2 (n = 666 cells) (Fig. S3B). These shape modes provide a meaningful and concise quantitative description of macrophage morphology, providing significant insights that may underly macrophage phenotypes and functionalities depending on their activation status.

### Morphological characterization of macrophage phenotypes

Macrophages are multifaceted, which is determined by the surrounding microenvironment, as well as multifunctional depending on their activation status, which assumes a variety of cell shapes. BMDMs when cultured in vitro and stimulated with cytokines to induce M1 or M2 polarization, displayed markedly different cell morphologies [30]. While unstimulated M0 macrophages show a round or slightly stretching appearance, addition of LPS, which stimulate M1 polarization, cause cell spreading with multiple protrusions within just 24 h of stimulation. In contrast, addition of IL-4, which stimulate M2 polarization, led to cell elongation (Fig. S4A). Quantitative analysis using Celltool [31] confirmed these three orthogonal modes of shape variability, which account for 86% of the total variation in shapes observed (see Supplementary information). The existence of only a few meaningful modes in combined macrophage populations implies that the phase space which macrophages reside is in a relatively limited subregion of the space of all possible shapes. In the literature, researchers have used various metrics to describe cell shape [31, 32]. There are several cell shape metrics like the cell area, perimeter length, convex area, major/minor-axis length, and many other. The best choice of metrics depends on the class of cell shapes that are found in the dataset. Here we aim to characterize and quantify the three primary shape modes identified above —circular (mode 1), “with protrusions” shape (mode 2), and elongated (mode 3) through clearly defined morphological parameters. Different combinations and number of parameters were explored. We utilized the following three parameters to characterize the morphological features of macrophage phenotypes, particularly given their uneven boundaries: compactness - to identify the complex and irregular boundaries of the cells; eccentricity - to quantify cell elongation; and solidity - to determine cell density. The combination of compactness, eccentricity, and solidity provided valuable and independent information to describe and differentiate the three primary shape modes identified within the combined macrophage populations.

In Fig. S4, we present examples of shape parameter values for the three major cell shapes, showcasing low, medium, and high parameter values for each shape. These parameter values effectively differentiate between the three primary shape modes and demonstrate the sufficiency of their combination for shape identification. Fig. 2 describes and defines these parameters calculated based on the number of pixels in a cell, its boundaries, the distances between them, and the cell trajectories. For the morphological characterization of macrophage subtypes, we take the unsupervised learning approach where the cells are not labeled as M0, M1, and M2 from their image names. We perform k-means clustering [33] with an optimal k value of 4 derived using silhouette score [34] on the three morphological parameters compactness, eccentricity, and solidity to understand the shape of the macrophages. The silhouette coefficient score plot and the silhouette plots for various k-values are shown in (Fig. S5). Fig. 3A shows the cluster assignment for the cells within the three-dimensional space of their morphological parameters’ compactness, eccentricity, and solidity. The identified 1018 cells from parent images M0, M1 and M2 are distributed into four clusters. The following observations are made about each cluster where the data points are represented as swarm plots [35, 36] (Fig. 3B-D):

**Fig 2.**
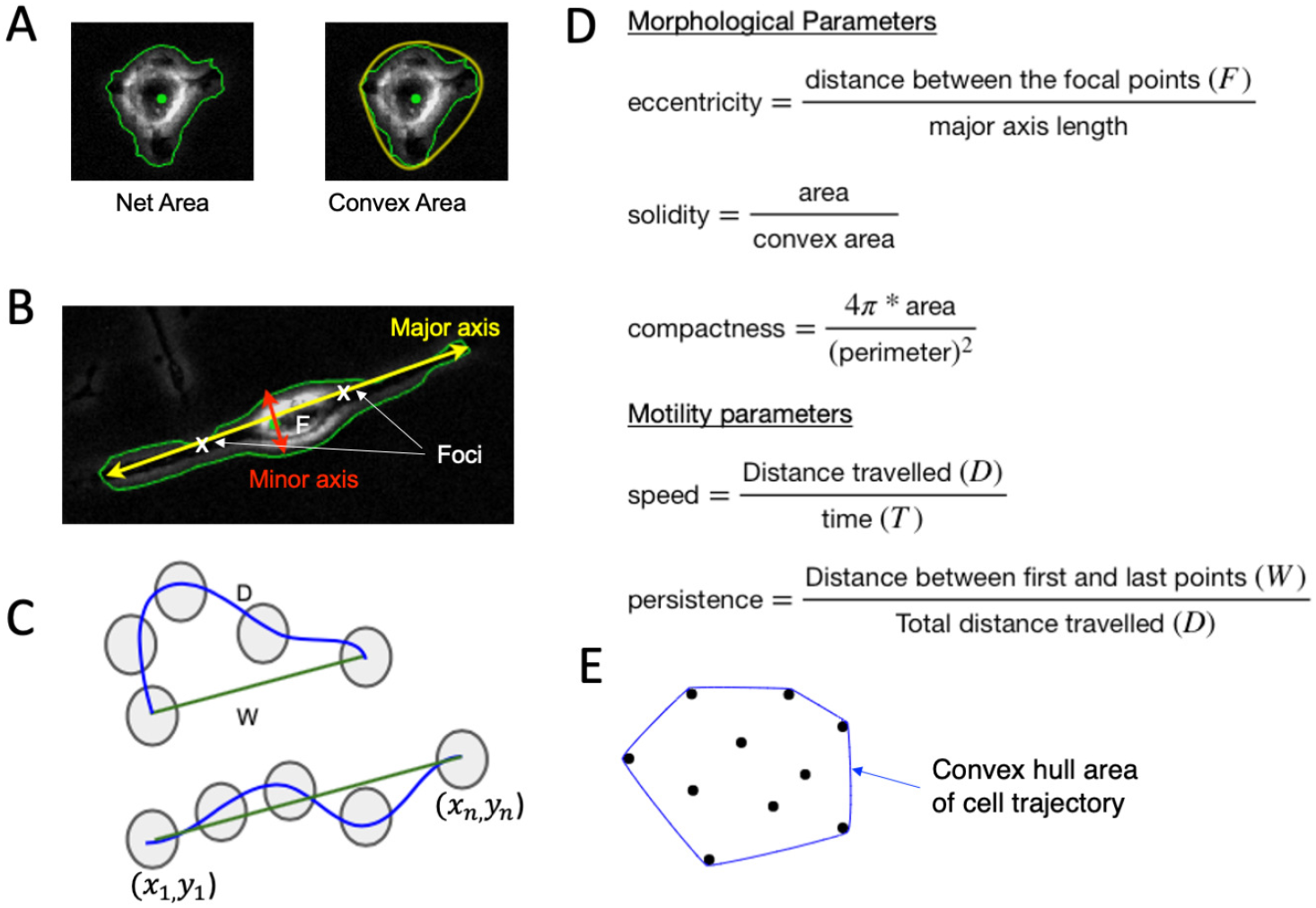
Defining the morphological and motility parameters. Morphology is defined by metrics related to the (A) area and perimeter of a cell and its convex hull, in addition to the (B) major and minor axis, while motility patterns are defined by (C) displacement metrics, (D) the corresponding formulas are used for morphological/motility parameters and (E) convex hull area of the cell trajectory points.

**Fig 3.**
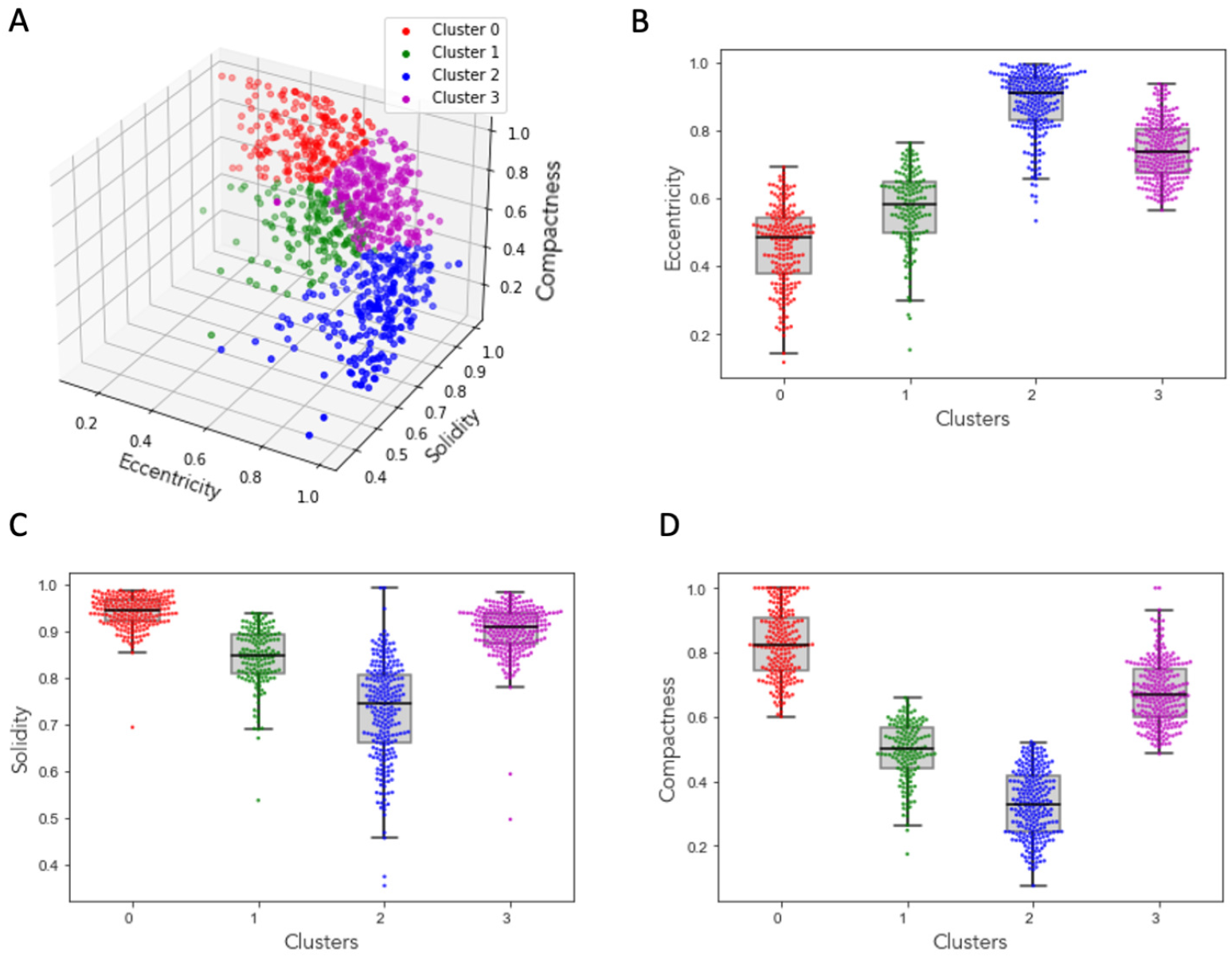
Clustering macrophages based on morphological parameters. (A) 3D plot of morphological parameters eccentricity, solidity and compactness of the cells showing four distinct clusters. Box plots with overlayed scatter points of eccentricity (B), solidity (C), and compactness (D).

Cluster 0: The cluster of cells with the lowest eccentricity values, highest compactness values, and highest solidity values indicates that the cells in this group are mostly circular, have smooth boundaries with little or no irregular boundaries, and are dense (regular boundaries without any visible holes within the cell structure). Fig. S4C shows an example cell from this cluster, along with their corresponding eccentricity, solidity, and compactness values.

Cluster 1: The cluster of cells with the lowest eccentricity values, lowest compactness values, and lowest solidity values suggests that the cells in this group are closer to being circular in shape, have highly irregular boundaries, and lack density, potentially due to their irregular boundaries or the presence of holes within the cell. Fig. S4B shows an example cell from this cluster, along with their corresponding eccentricity, solidity, and compactness values.

Cluster 2: The cluster of cells with higher eccentricity values, above-average compactness values, and higher solidity values suggests that the cells in this group are closer to being elliptical in shape, have relatively smoother boundaries, and are dense (regular boundaries without any visible holes within the cell structure). Fig. S4A shows an example cell from this cluster, along with their corresponding eccentricity, solidity, and compactness values.

Cluster 3: The cluster of cells with higher eccentricity values, lowest compactness values, and solidity values ranging from least to average suggests that the cells in this group are elliptical in shape, have highly irregular boundaries, and are not densely packed. In Fig. S7, we present examples of cells from each cluster. Cluster 3 contains 333 cells, with 28% from M0, 41% from M1, and 31% from M2 images. Our analysis revealed that this cluster includes indifferent groupings, cell overlapping, or transitional cells. The shape of these cells does not align with any of the primary modes identified in the principal component analysis.

From the above observations, we can describe cells in Cluster 0 as circular cells, cells in Cluster 1 as the cells with protrusions, and cells in Cluster 2 as the elongated cells.

### Motility patterns can be used to differentiate cell morphology-based groups

We would like to show that the chosen displacement metrics of speed and persistence can be used to differentiate cell groups based on morphology. To do this we apply discriminant analysis, a multivariate technique, to separate two morphology-based groups based on measured speed and persistence. This helps to identify the contribution of each variable in separating the groups. Here, we show that a cell’s morphological features are in fact linked closely to its motile behavior. This relationship exists independent of the macrophage subtype. For this we leverage each of the unique cells identified above in clusters and quantify their corresponding motility patterns. To investigate this relationship, we identify a minimum set of motility parameters that capture the set of observed migratory behavioral patterns. We compute speed and persistence (a measure to find how far a cell has moved from its initial time step) for all the cells in clusters 0-2 (Fig. 4A).

**Fig 4.**
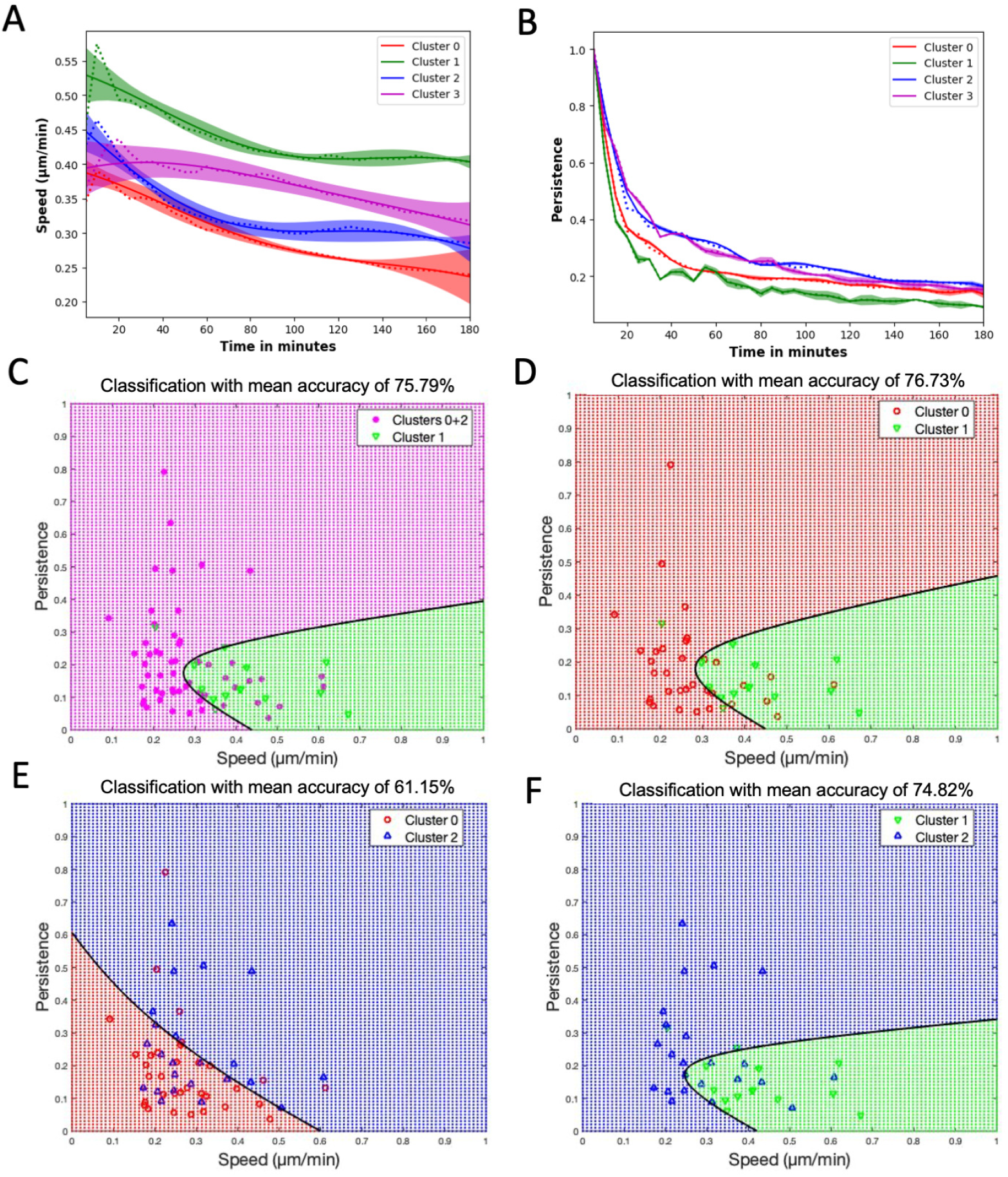
Classification and discrimination of morphological groups using motility parameters. (A) Speed and (B) Persistence, where dotted lines are the mean values of the corresponding clusters, solid lines are the Gaussian Process Regressor model (GPR) predictions, and the color bands are their 95% confidence intervals. (C) Plot showing the quadratic classification of the cells using Speed vs Persistence between Cluster 1 cells and combined clusters 0 and 2 cells with a classification mean accuracy from 5-fold cross-validation of around 76%. (D) Plot showing the quadratic classification of the cells using Speed vs Persistence between Cluster 0 cells and Cluster 1 cells with a classification mean accuracy of around 77%. (E) Plot showing the quadratic classification of the cells using Speed vs Persistence between Cluster 0 cells and Cluster 2 cells with a classification mean accuracy of around 61%. (F) Plot showing quadratic the classification of the cells using Speed vs Persistence between Cluster 1 cells and Cluster 2 cells with a classification mean accuracy of around 75%. The black line in C-F is the decision boundary between the classes. Additionally, the classification mean accuracies of Clusters 1+2 vs Cluster 0 and Clusters 0 +1 vs Cluster 2 is around 55% and 53% respectively.

We use the Gaussian process regressor (GPR) model [32, 37] to plot the speed and persistence of the cells. The GPR model utilizes a Gaussian process to model the distribution of the data and predict outcomes based on the observed data points. In Fig. 4, each cluster’s results are depicted using a distinct color. The GPR model’s mean is represented by a solid line, while the average value computed from the data at each time point is indicated by dashed lines. Additionally, the band in the plot represents a 95% confidence interval. Fig. 4A indicates that the speed of cells in Cluster 1 (protruded cells) is the highest, followed by the speed of cells in Cluster 2 (elongated cells), and the lowest speed is observed for cells in Cluster 0 (circular cells). Fig. 4B displays that the cells in Cluster 2 exhibit the highest persistence, followed by Cluster 0, with the lowest persistence observed for the cells in Cluster 1. Combining the motility parameters analysis with morphological clustering analysis, we observe that circular cells move slower and stay close to their initial time frame position; protruded cells move faster but still stay close to their initial time frame position; whereas elongated cells move slower but reach farther away from their initial time frame position. For Cluster 3 cells, which exhibited indifferent cell groupings or overlapping cells, we can observe that their speed falls between Cluster 1 and the remaining clusters. Also, these cells display a high persistence level, as observed in Cluster 2 cells.

To better visualize and quantify the relationship between cell morphology and motility we perform quadratic discriminant analysis on the motility parameters speed and persistence for the clusters from the morphological clustering analysis. This analysis provides a binary classification based on two input features (i.e., speed and persistence). With this method we can determine to what extent we can predict the cells cluster of origin (e.g., predict morphological characteristics) based on its computed speed and persistence (Fig. 4, showing the results for various pairs of classification labels). We perform 5-fold cross-validation and calculate the mean accuracy. Classification accuracy percentages confirm that the Cluster 1(protruded cells) can be distinguished from the Cluster 0 (circular cells) and Cluster 2 (elongated cells) combined or independently using the speed and persistence motility parameters (Fig. 4C, D, F). However, the classification accuracy percentage between Cluster 0 (circular cells) and Cluster 2 (elongated cells) is comparatively lower around 61% (Fig. 4E). The ability to predict morphological features based on cell motility imply a strong relationship between the two.

### Motility characterization can be used to predict macrophage subtypes

We have shown that macrophage subtypes can be predominantly characterized by specific combinations of morphological features. Additionally, we have shown evidence to suggest that there is a relationship between motile behavior and morphology. Thus, we expect that macrophage subtypes should exhibit distinct motile behavior. To demonstrate that motility can be used as a substitute for morphological features, we quantify motile behavior for each cell from the cell tracking results by computing speed and persistence for each macrophage subtype given by the parent image label and taking a supervised learning approach. Based on this motility parameter analysis given in (Fig. S6A, B), we find that M0 cells move slower and stay close to their initial time frame position; M1 cells move faster and but still stay close to their initial time frame position; whereas M2 cells move slower and reach farther away from their initial time frame position. These motility observations can act as an independent model in the identification of the macrophage subtypes without analyzing the morphology of the cells. Quadratic discriminant analysis is also performed (Fig. S6C-F) to quantify the motility of the cells in M0, M1 and M2 images. We note that the mean accuracies are the same or worse when compared to those using cluster labels due to the heterogeneity observed for each parent label based on cell culturing protocols. Recall that the three orthogonal modes of shape variability account for 86% of the total variation in shapes observed. Furthermore, approximately 30% of the cells were placed into Cluster 3, which included cells with mixed or ambiguous features. Thus, a classification accuracy of around 70% should be considered formidable when using only morphology. Below we investigate accuracies achieved with motility information alone and combined information, respectively.

### Increased prediction accuracy of macrophage subtype is obtained using both morphological and motility features

We note that in the previous analysis some macrophage subtypes are more easily distinguished by motility parameters and others by morphological features. Thus, we expect that combining both morphological and motile behavior can improve our ability to classify macrophage subtypes. The advantage of this would be an ability to classify macrophage subtypes using only phase contrast images. Additionally, we present a generalizable machine learning model to properly compare predictive capabilities based on both motility and morphology independently and combined. We compute eccentricity, compactness, and cell size to observe morphological features, and convex hull area of cell trajectory, speed, and persistence to extract motility features for all cells in M0, M1, and M2 images (Fig. 5). For our machine learning classifier, we consider a support vector machine (SVM) [38] model, which can deliver high precision and accuracy regardless of the number of attributes and data instances [35]. We have 1018 cells from all the image sets where 251 are from M0 images, 369 from M1 images and 398 from M2 images. We shuffle the data and use stratified 5-fold cross-validation. During each iteration, the dataset was divided into approximately [200, 296, 318] cells for M0, M1, and M2 classes in training and [73, 80, 51] cells for M0, M1, and M2 classes in testing sets. We now use Synthetic Minority Oversampling Technique (SMOTE) (35) to balance the minority classes M0 and M1. The final training dataset has 954 datapoints, with 318 datapoints from each of the M0, M1, and M2 image classes. We fit the training data using SVM classifier -radial basis function (RBF) kernel and predict the output on the test data where the output of the model is the macrophage subtype given by the parent image label. Fig. 5A shows the schematic of the machine learning model with the features, input labels and the expected output. To demonstrate that the combination of morphology and motility enhances the classifier prediction, we develop three models. The first model is trained using only morphological features (eccentricity, compactness, cell size) as inputs. The second model is trained using only motility features (convex hull area of cell trajectory, speed, persistence) as inputs. The third model is trained using both morphological and motility features as inputs. The results are shown in Fig. 5B-D, where the prediction accuracy of the model on test data is plotted in a confusion matrix, where rows represent the actual labels and columns represent the predicted labels. The morphology trained model showed that the prediction accuracies for M0, M1 and M2 labelled cells are 52%, 69% and 70% respectively (Fig. 5B). While the motility trained model had the prediction accuracies for M0 (48%) and M1 (68%), M2 cells (71%) (Fig. 5C). Strikingly, the combined morphology and motility trained model significantly improved prediction accuracies for all the M0, M1 and M2 cells to 60%, 72% and 74% respectively (Fig. 5D). Hence, by using the combined morphological and motility parameters we can predict the macrophage subtypes with an improved accuracy compared to the models which use either morphology or motility alone (Table S1). For reference, when analyzing individual biomarkers in flow cytometry, we note that achieving a prediction accuracy of 74% is considered high, owing to variability in biomarker expression influenced by experimental context [39]. Thus, the prediction accuracies obtained are meaningful within a biological context and support the claim that there exists a relationship between cell motility, cell morphology, and macrophage subtypes.

**Fig 5.**
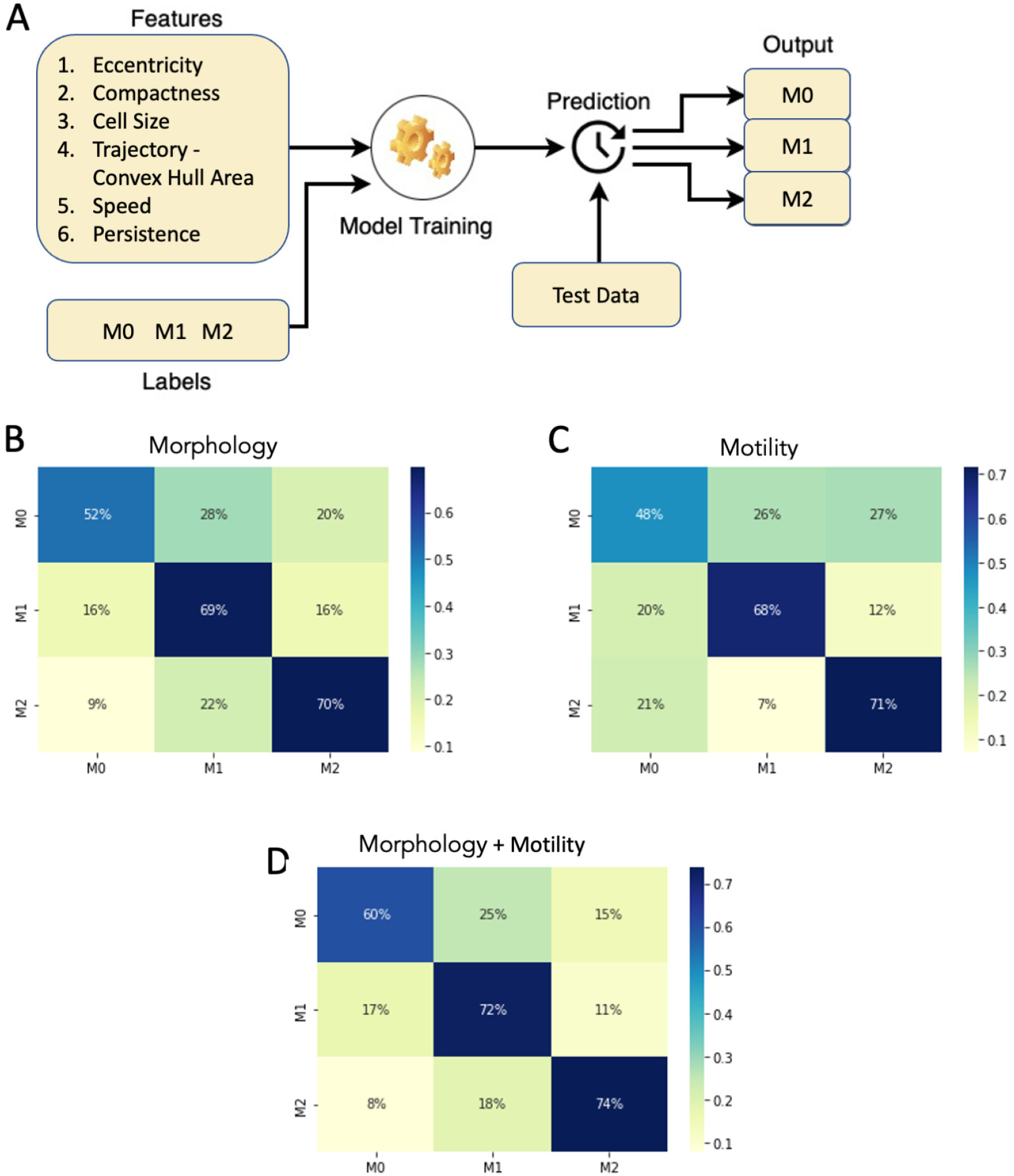
Classification of macrophage subtypes using SVM on morphological and motility parameters - Stratified 5-fold cross-validation plots for the SVM model. (A) Schematics of the input and output of the machine learning model. SVM classification results showing the percentage distribution of the cells in a confusion matrix with columns being the predicted labels and rows being the actual labels. (B) Input features with morphological parameters (Eccentricity, Compactness, Cell Size). (C) Input features with motility parameters (Convex Hull Area of the trajectory, Speed, Persistence). (D) Input features with both morphological and motility parameters.

## Discussion

It is well known that macrophages cannot be cleanly grouped into their subtypes, and in fact, various intermediate subtypes exist, yet their artificial classification is important to understanding gene regulatory pathways involved in the various functions carried out by the full spectrum of macrophage phenotypes [40]. Such understandings can help establish ways to target and control their activity to improve health outcomes in applications such as wound healing.

Image-based methods provide a fast and efficient way to classify macrophage subtypes. Previous work has focused on machine learning methods for predicting macrophage subtypes based on morphology [30, 41]. Machine learning effectively substitutes the manual classification of cell phenotype by researchers and clinical practitioners and offloads the decision-making to an algorithm. The reasons automated image processing, segmentation, and analysis have found such interest are that they offer huge possibilities to free up researchers’ time and to ensure consistent interpretation of results free from any human bias. Prediction methods that rely on cell morphology require high-quality images to be able to accurately segment cells and characterize their morphology. In this work, we show that spontaneous cell motility patterns can also serve as unique signatures for macrophage subtypes. Furthermore, an impressive amount of cell tracking algorithms [42], benchmark techniques [43, 44] and software [13–15, 45, 46] have been developed and made available to the scientific community which are more ubiquitous and robust to blurry images. However, we note that it can be a complex task to extract the morphology of a cell if it were to be suspended rather than spontaneously moving on a substrate. We also recognize that the motility can be affected by the substrate itself [47]. This means that changing the substrate may require new characterization for predictions. However, this may imply that the different cell types move differently in different environments. We know the extracellular matrix changes over time in wound healing, which allows different cell types to carry out their work.

Independently, motility and morphological features can provide ambiguous discriminant information depending on the macrophage subtype. For example, the prediction accuracies derived from motility-based information closely match those obtained through morphology-based predictions. Thus, combining both cell motility and morphological information can reliably and independently detect non-activated naïve macrophages, or macrophages activated by either lipopolysaccharide (LPS) or interleukin 4 (IL-4) exposure at single-cell level. Furthermore, the numerous metrics applied to characterize the morphology and migration patterns are simplified representations (e.g. information is lost). One may consider exploring ML methods to extract patterns with nuanced information from the raw data.

We note that the images used for analysis required that cell densities remain low. We noted that some of the cells in cluster 3 could likely be clusters of cells that are identified as a single cell due to the segmentation regions formed when the cells are very close to each other. Further improvement of the segmentation algorithm to correctly identify the high-density cell regions and post-processing methods to remove the cells for analysis beyond a particular cell area, might lead some of these cells to get distributed in the above three clusters. Additionally, Cluster 3 also seemed to include cells that had combined features from two or more of the remaining clusters. Thus, Cluster 3 may include macrophages that are in a transient state between subtypes or may represent an ambiguous state not yet explored. The methods in this paper could potentially be used to further understand the full spectrum of macrophages and track potential shifts in macrophage subtypes over time. Furthermore, a future adaptation of this method can be used in bioengineering applications to characterize and quickly screen genetic variants such as in engineered CAR-Macrophages [48].

## Conclusion

In this paper, we present that using image processing techniques, we can characterize cell behavior and migratory patterns that can be fed into our machine-learning model to predict macrophage subtypes. While we recognize that other methods such as flow-cytometry can offer improved accuracy in macrophage classification [39],our method has the advantage of being non-invasive. It does not require cell fixation or labeling, preserving cell viability for further functional assays. Additionally, this approach allows for real-time analysis. The ability to assess macrophage polarization through dynamic migration and morphology changes offers insights into the real-time behavior of cells in response to various stimuli. Our method also has the potential for high-throughput screening. Once fully optimized, it could facilitate high-throughput screening of biochemical effects on macrophage polarization, providing a valuable tool for both basic research and drug discovery. Analysis of these dynamics without any external cues given like electrical guidance and staining would give us insights that are critical to quantifying macrophage recruitment and activation in vivo. In future work, this analysis could also be applied to the images in galvanotaxis to observe the dynamics of the cell behavior with and without electric fields. Furthermore, a direct comparison between fixed tissue analysis and flow cytometry, could enhance understanding of the potential applications of our approach. While this comparison was not conducted in the current study due to its exploratory nature and focus on developing the machine learning model, this is a potential direction for future research. Finally, this method can be adapted to serve other cell types and applications but may require identifying more suitable morphological and displacement metrics where applicable.

## Data Availability

The complete dataset is available at https://datadryad.org/stash/share/vrQi68_C27sbdkP2uAXUE8kvy_BxlYkRImI5f6_9WY0 with a metadata file accompanying the dataset. The source code required to reproduce the findings of this study has been uploaded to the following GitHub repository: https://github.com/Gomez-Lab/CellAnalysis.

