## Supplemental Table 1 for "A data-driven approach to establishing cell motility patterns as predictors of macrophage subtypes and their relation to cell morphology"

**Table S1. Macrophage subtype classification with accuracy percentages for different analysis techniques used in the study**

| Input parameters | Method | M0 | M1 | M2 |
|---|---|---|---|---|
| Morphology | K-means Clustering | 38% | 52% | 46% |
| Morphology | SVM | 69% | 71% | 71% |
| Motility | SVM | 75% | 75% | 41% |
| **Morphology + Motility** | SVM | **81%** | **79%** | **79%** |
