## Supplementary figures and images for "A data-driven approach to establishing cell motility patterns as predictors of macrophage subtypes and their relation to cell morphology"

### Supplemental Fig 1

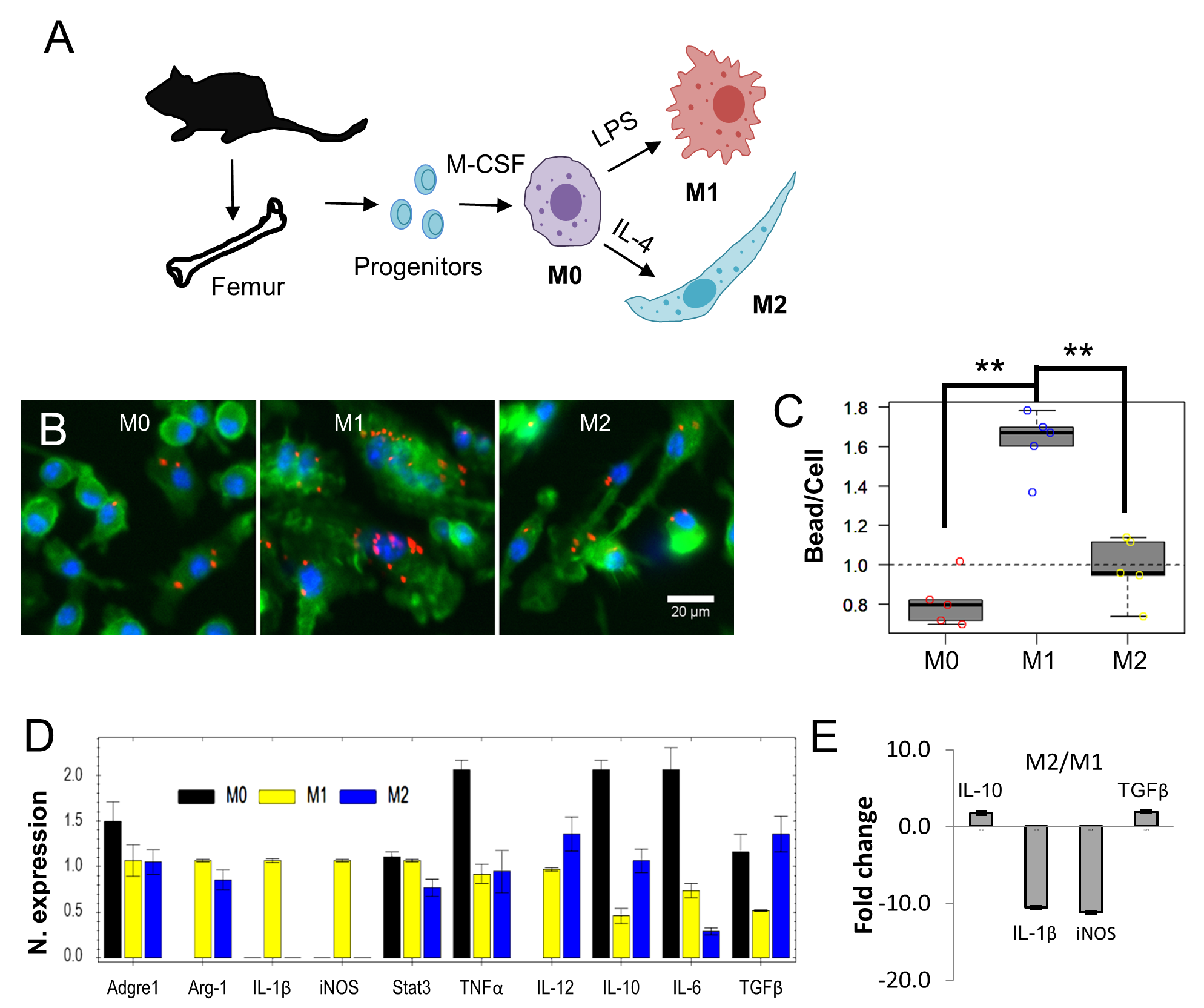

### Supplemental Fig 2

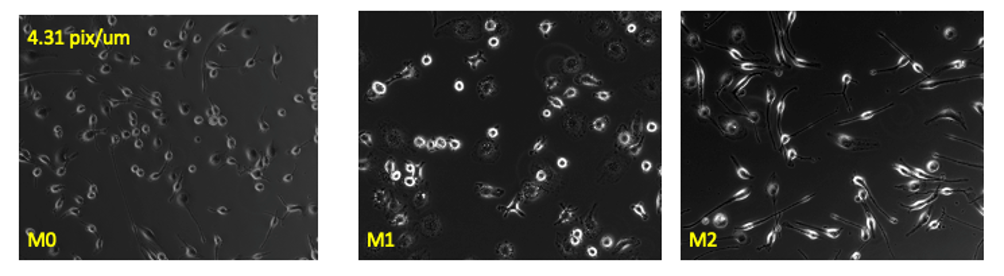

### Supplemental Fig 3

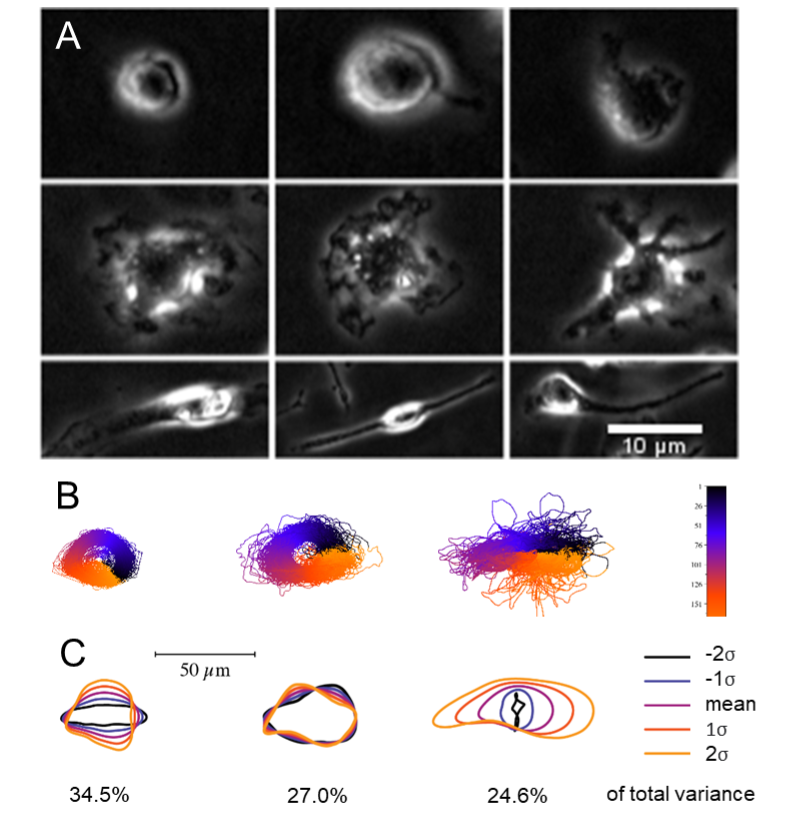

### Supplemental Fig 4

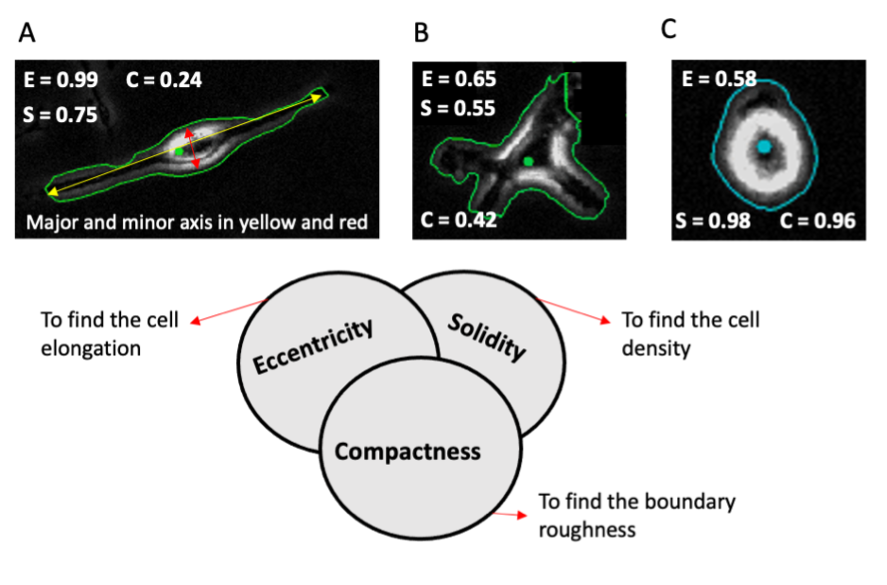

### Supplemental Fig 5

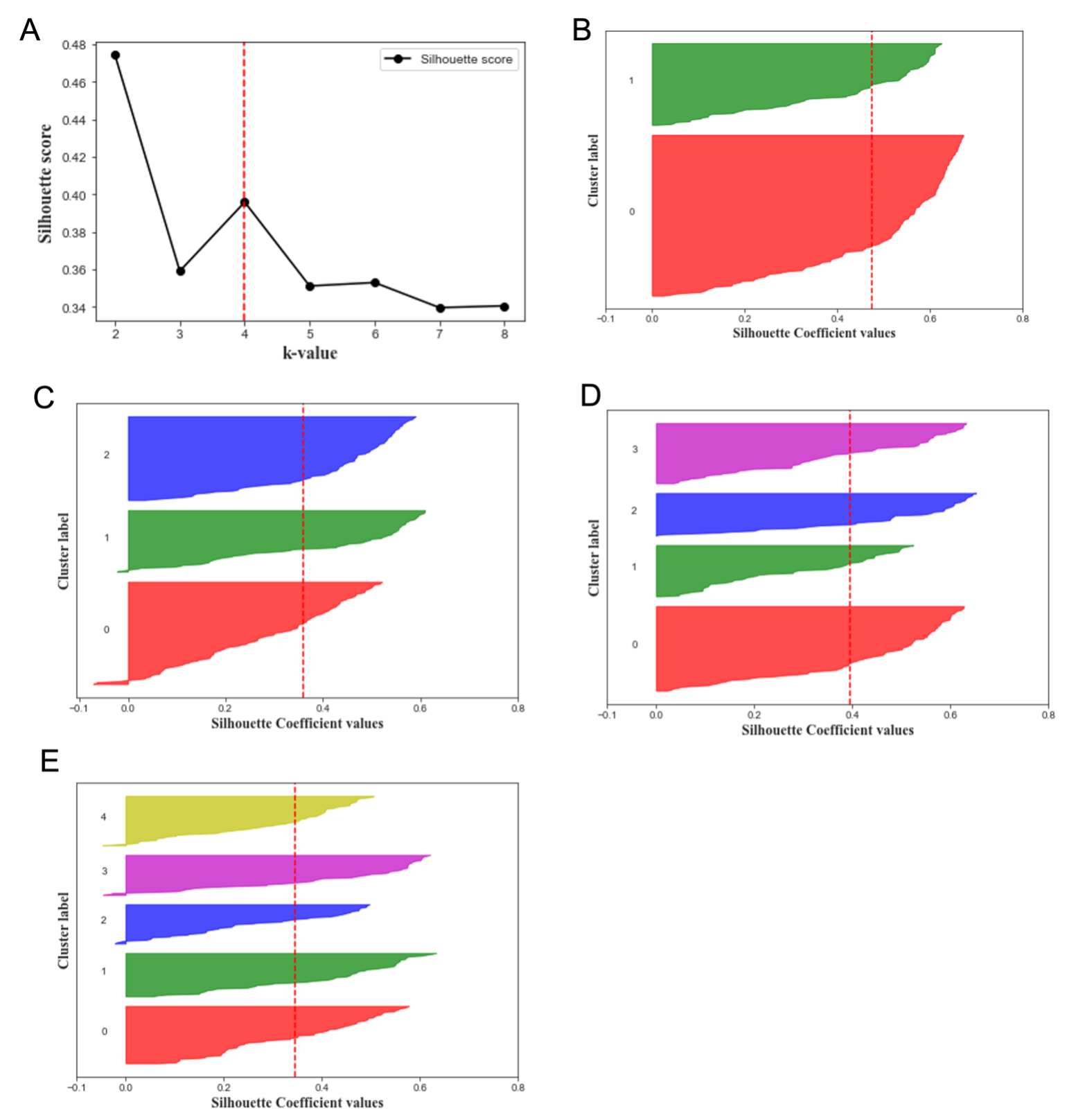

### Supplemental Fig 6

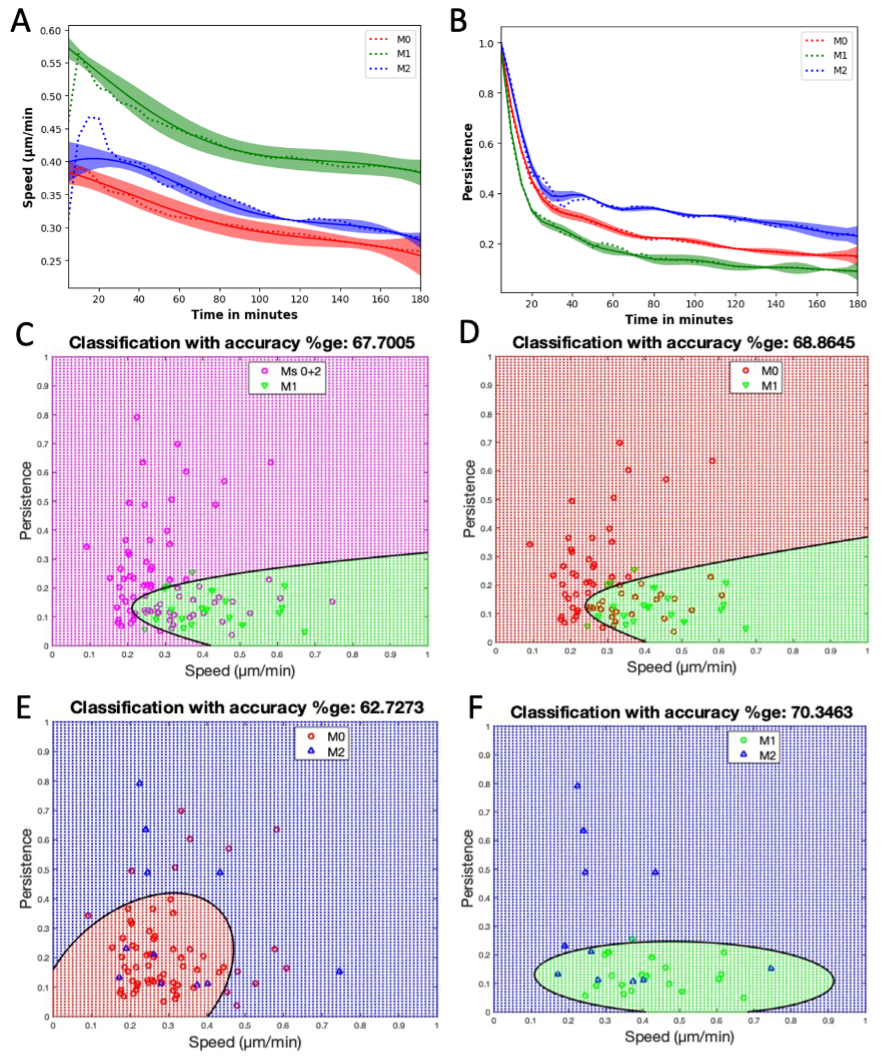

### Supplemental Fig 7

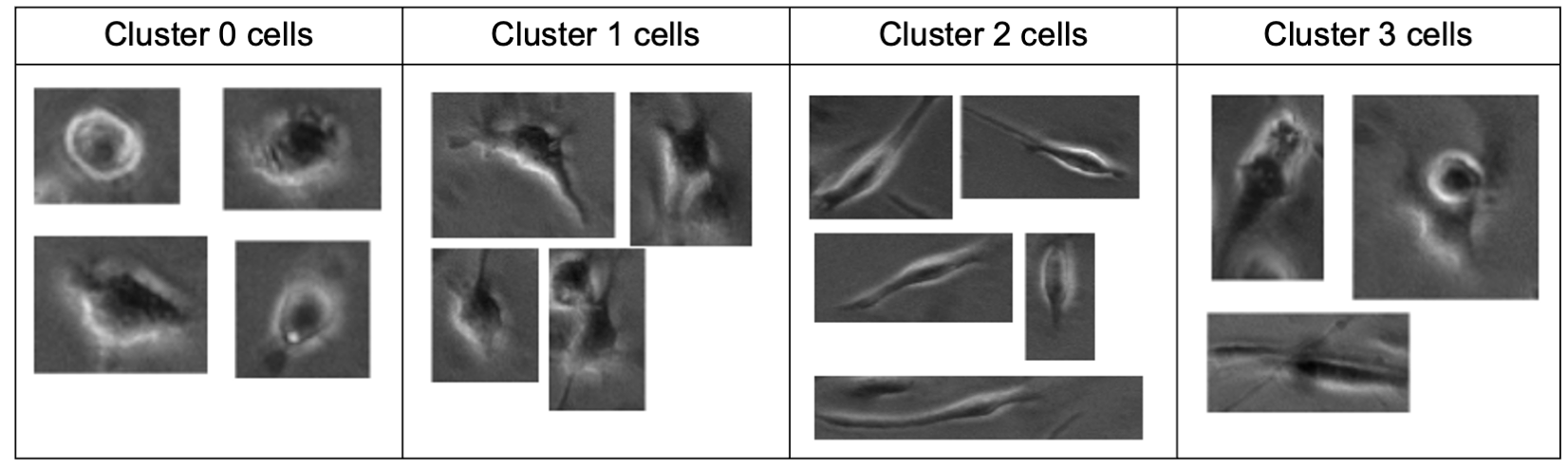
