## Supplemental Table 2 for "A data-driven approach to establishing cell motility patterns as predictors of macrophage subtypes and their relation to cell morphology"

**Table S2: Target genes and primers used in q-PCR.**

| Target gene | Forward primers (5'-3') | Reverse primers (5'-3') | Macrophage Subtypes |
|---|---|---|---|
| Arg-1 | CGTTGTATGATGCACAGCCG | CCCCACCCAGTGATCTTGAC | M1 |
| CD206 | GTTCACCTGGAGTGATGGTTCTC | AGGACATGCCAGGGTCACCTTT | M2 |
| CD31 | ACAGAGCCAGCAGTATGA | AATGACAACCACCGCAAT | Angiogenesis |
| CXCL10 | GATGGATGGACAGCAGAG | GGAAGATGGTGGTTAAGTTC | M1 |
| Adgre1 | TCTGGGGAGCTTACGATGGA | GAATCCCGCAATGATGGCAC | Macrophage |
| Fizz-1 | CCTGCTGGGATGACTGCTACT | AGATCCACAGGCAAAGCCAC | M2 |
| GAPDH | TCTCCTGCGACTTCAACA | TGTAGCCGTATTCATTGTCA | Housekeeping |
| IL-10 | CAGAGCCACATGCTCCTAGA | TGTCCAGCTGGTCCTTTGTT | M2 |
| IL-12p40 | TGGTTTGCCATCGTTTTGCTG | ACAGGTGAGGTTCACTGTTTCT | M1 |
| IL-1β | GTGCAAGTGTCTGAAGCAGC | CAAAGGTTTGGAAGCAGCCC | M1 |
| IL-6 | GGAGTCACAGAAGGAGTGGC | CGCACTAGGTTTGCCGAGTA | M2 |
| iNOS | GTTCTCAGGCCAACAATACAAGA | GTGGACGGGTCGATGTCAC | M1 |
| MCP-1 | AGCCAACTCTCACTGAAGCC | GGACCCATTCCTTCTTGGGG | M1 |
| MMP-9 | CTGGACAGCCAGACACTAAAG | CTCGCGGCAAGTCTTCAGAG | M2 MMP |
| PPAR-γ | TCCTGTAAAAGCCCGGAGTAT | GCTCTGGTAGGGGCAGTGA | M2 Metabolic |
| Stat3 | CTTGTCTACCTCTACCCCGACAT | GATCCATGTCAAACGTGAGCG | M1 TF |
| Stat6 | TGAGGTGGGGACCAGCCGG | GTGACCAGGACACACAGCGG | M2 TF |
| TGF-β1 | TGGAGCAACATGTGGAACTC | CAGCAGCCGGTTACCAAG | M2 |
| TLR4 | CAGAACAATAGAAGAGGAAGAC | GGCACTAACCACATAGAGAA | M1 |
| TNF-α | GACGTGGAACTGGCAGAAGA | ACTGATGAGAGGGAGGCCAT | M1 |
| Ym-1 | CTCACTTCCACAGGAGCAGG | AGCTGCTCCATGGTCCTTC | M2 |
| β-actin | GGCTGTATTCCCCTCCATCG | CCAGTTGGTAACAATGCCATGT | Housekeeping |
